# Somatic genome editing with the RCAS/TVA-CRISPR/Cas9 system for precision tumor modeling

**DOI:** 10.1101/162669

**Authors:** Barbara Oldrini, Álvaro Curiel-García, Carolina Marques, Veronica Matia, Özge Uluçkan, Raul Torres-Ruiz, Sandra Rodriguez-Perales, Jason T. Huse, Massimo Squatrito

## Abstract

It has been gradually established that the vast majority of human tumors are extraordinarily heterogeneous at a genetic level. To accurately recapitulate this complexity, it is now evident that *in vivo* animal models of cancers will require to recreate not just a handful of simple genetic alterations, but possibly dozens and increasingly intricate. Here, we have combined the RCAS/TVA system with the CRISPR/Cas9 genome editing tools for precise modeling of human tumors. We show that somatic deletion in neural stem cells (NSCs) of a variety of known tumor suppressor genes (*Trp53*, *Cdkn2a* and *Pten*), in combination with the expression of an oncogene driver, leads to high-grade glioma formation. Moreover, by simultaneous delivery of pairs of guide RNAs (gRNAs) we generated different gene fusions, either by chromosomal deletion (*Bcan*-*Ntrk1)* or by chromosomal translocation (*Myb-Qk*), and we show that they have transforming potential *in vitro* and *in vivo*. Lastly, using homology-directed-repair (HDR), we also produced tumors carrying the Braf V600E mutation, frequently identified in a variety of subtypes of gliomas. In summary, we have developed an extremely powerful and versatile mouse model for *in vivo* somatic genome editing, that will elicit the generation of more accurate cancer models particularly appropriate for pre-clinical testing.

## Introduction

A decade of studies has underlined the complexity of the genetic events that characterize the genomic landscapes of common forms of human cancer ^1^. While a few cancer genes are mutated at high frequencies (>20%), the greatest number of cancer genes in most patients appear at intermediate frequencies (2–20%) or lower ^2^. Strikingly, the functional significance of the vast majority of these alterations still remains elusive. A current high priority in cancer research is to functionally validate candidate genetic alterations to distinguish those that are significant for cancer progression and treatment response. In order to do so, it is essential to develop flexible genetically engineered mouse models that can speed up the functional identification of cancer driver genes among the large number of passenger alterations ^3^.

The growing level of sophistication of the genome engineering technologies has made it possible to target almost any candidate gene in the *in vivo* setting. The CRISPR (<u>C</u>lustered <u>R</u>egularly <u>I</u>nterspaced <u>S</u>hort <u>P</u>alindromic repeats) – Cas (<u>C</u>RISPR-<u>a</u>ssociated), the most powerful genome editing system so far, has revolutionized research in many fields, including cancer animal modelling, by allowing precise manipulation of the genome of individual cells. Its applications span from the inactivation of tumor suppressor genes, to the generation of somatic point mutations and more complex genomic rearrangements such as gene fusion events.

A possibly significant limitation of CRISPR-based *in vivo* somatic genome editing is the requirement to concurrently deliver the RNA guides and the Cas9 enzyme to the specific tissue of interest. To deal with this issue, various groups recently generated transgenic mice expressing Cas9 in a Cre- or tetracycline-dependent manner ^4–6^. The combination of somatic genome editing with the vast collection of currently available genetically engineered mouse models will provide the chance to introduce defined genetic lesions into specific cell types, leading to the development of more accurate tumor models.

The RCAS/TVA based approach uses replication-competent avian leukosis virus (ALV) splice-acceptor (RCAS) vectors to target gene expression to specific cell types in transgenic mice. In these mice, cell type-specific gene promoter drive expression of TVA, the cell surface receptor for the virus. The RCAS/TVA system has been successfully used in different mouse models to deliver genes or shRNAs of interest into a plethora of cell types: neural stem cells, astrocytes, hepatocytes and pancreatic acinar cells, among many others ^7^.

Here we describe a series of new mouse models that combine the genome editing capability of the CRISPR/Cas9 system with the somatic gene delivery of the RCAS/TVA approach to generate precision tumor modeling. To prove the efficacy of such a powerful system, we produced a number of *in vivo* and *ex vivo* models of glioma with tailored genetic alterations.

The gliomas are a large group of brain tumors and within gliomas the glioblastoma (GBM) is the most frequent form of the disease and overall the most common and lethal primary central nervous system (CNS) tumor in adults. A series of large-scale genomic analysis has underlined the complexity of the genetic events that characterize the glioma genome. However, so far, we have been able to study only a minority of these genetic alterations due to the lack of appropriate tumor models.

We took advantage of the previously developed *Rosa26-LSL-Cas9* (*LSL-Cas9*) knockin mouse strain ^4^ and combined it with the *Nestin-tv-a* (*Ntv-a*) and the *GFAP-tv-a* (*Gtv-a*) transgenic mice that express the TVA receptor under the control of the rat *nestin* and human *GFAP* promoter, respectively ^8,9^. Moreover, we have further crossed those strains with either the *Nestin-Cre* (*Nes-Cre*) or *hGFAP-Cre* transgenic lines ^10,11^, to allow for Cas9 expression in neural stem/progenitor cells and in astrocytes, or with a mouse strain carrying a tamoxifen-activated recombinase, *hUBC-CreERT2* ^12^, for ubiquitous and inducible Cas9 expression.

Through *in vivo* delivery of RCAS plasmids that carry guided RNAs (gRNAs) for a series of tumor suppressor genes known to be frequently deleted in GBM (*Trp53*, *Cdkn2a* and *Pten*), in combination with the expression of the Platelet Derived Growth Factor Subunit B (PDGFB), we show that we can efficiently generate high-grade gliomas in mice that express both Cas9 and TVA in the Nestin or GFAP positive cells. Moreover, by simultaneous *ex vivo* transduction into neural stem cells (NSCs) of RCAS plasmids expressing pairs of gRNAs we generated either chromosomal deletion (*Bcan-Ntrk1* gene fusion) or chromosomal translocation (*Myb-Qk* gene fusion) and we show that they lead to glioma formation when transplanted in immunocompromised mice. We further generated *Braf* mutant gliomas by inducing a homology-directed repair-mediated BRAF V600E mutation.

Lastly, by *ex vivo* treatment of some of these tumor models we demonstrate their utility for pre-clinical testing of targeted therapies.

In conclusion, combining the RCAS/TVA and CRISPR/Cas9 models we have developed an extremely powerful mouse model for *in vivo* somatic genome editing, that allows targeting specific cell types with definite genetic alterations to generate precision tumor models.

## Results

### Generation of CNS Cas9-expressing mouse strains

To test the possibility of somatic genome editing by combining the RCAS/TVA and CRISPR/Cas9 models, we generated a series of mouse strains that allowed the TVA and Cas9 expression in specific cell types in the brain.

Nestin is an intermediate filament protein (IFP) that is predominantly expressed in the central nervous system stem/ progenitor cells during embryonic development, but also in muscles and other tissues. In adult organism, its expression in the brain is mainly restricted to the neural stem cell compartment of the subventricular zone (SVZ). After differentiation, nestin is downregulated and replaced by tissue-specific IFPs. The glial fibrillary acidic protein (GFAP) is an IFP that is expressed by numerous cell types of the CNS including astrocytes and ependymal cells. *Nestin-tv-a* (*Ntv-a*) and *GFAP-tv-a* (*Gtv-a*) transgenic mice that express the TVA receptor under the control of the rat *nestin* and human *GFAP* promoter, have been widely used for modeling brain tumorigenesis ^8,9,13^.

The Rosa26-LSL-Cas9 knockin mice (*LSL-Cas9*) have a floxed-STOP cassette precluding expression of the downstream bicistronic sequences (Cas9-P2A-EGFP) and it was generated to overcome the delivery challenges of the Cas9 enzyme to specific tissues of interest ^4^. We crossed these mice with the *Ntv-a* and *Gtv-a* transgenic mice to obtain the *Ntv-a; LSL-Cas9* and *Gtv-a; LSL-Cas9*. Although RCAS-Cre expressing plasmids have been previously used in combination with different TVA expressing mice to allow tissue specific deletion of a variety of floxed alleles ^13–15^, to ensure a robust recombination in the CNS of the floxed-STOP cassette in the *Ntv-a; LSL-Cas9* and *Gtv-a; LSL-Cas9* we further crossed these mice with either the *Nestin-Cre* (*Nes-Cre*) or *hGFAP-Cre* transgenic lines ^10,11^. The resulting *Ntv-a; Nes-Cre; LSL-Cas9* and *Gtv-a; hGFAP-Cre; LSL-Cas9* mice presented no abnormalities in development and size (Supplementary Fig. 1a), were fertile and had normal litter sizes.

**Figure 1:**
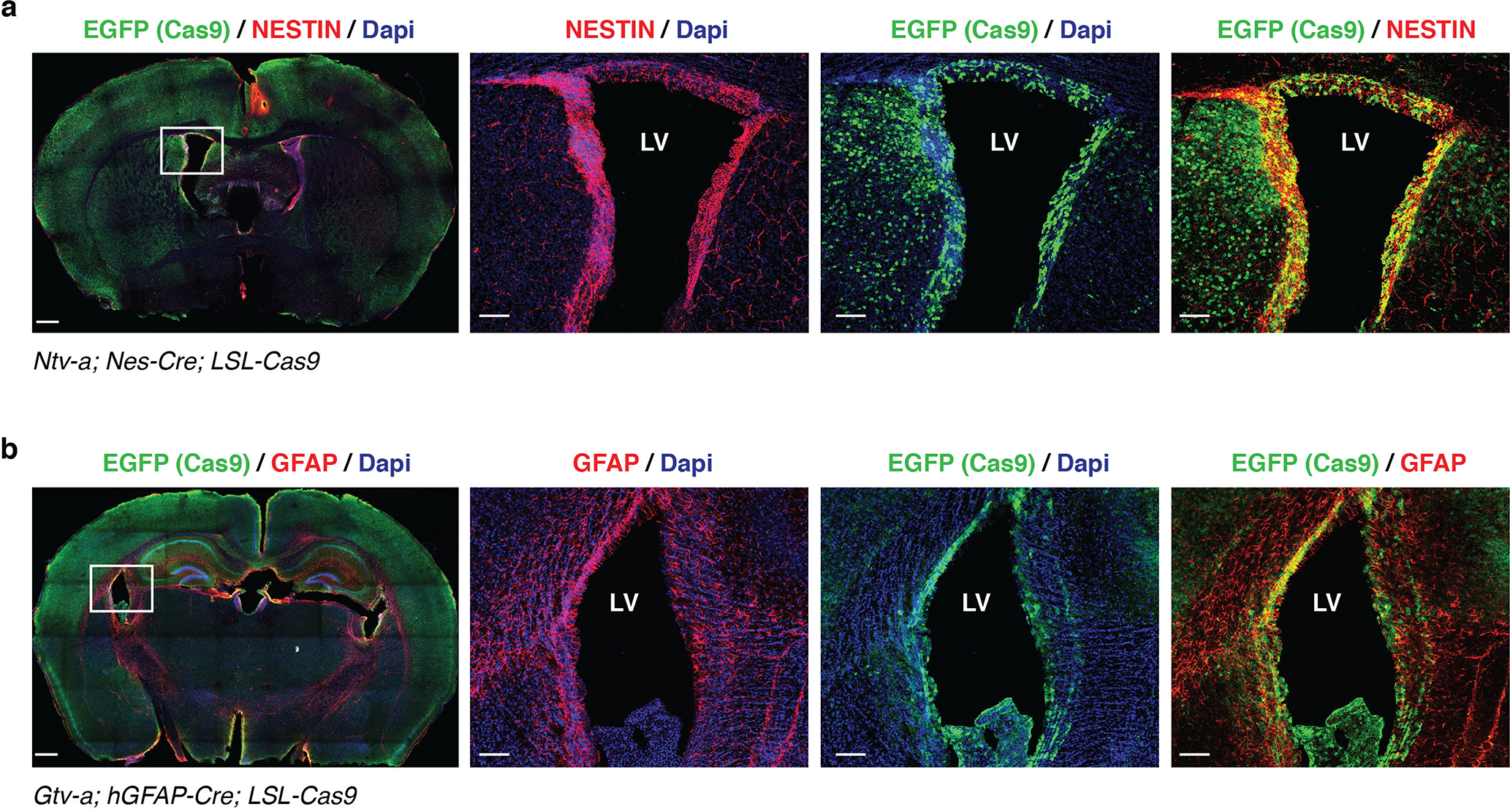
Cas9 expression in the brain of TVA/Cas9 mouse strains. **(a)** and **(b)** Immunofluorescence staining performed on brain sections of 4 weeks old *Ntv-a; Nes-Cre; LSL-Cas9* and *Gtv-a; hGFAP-Cre; LSL-Cas9* mice with antibody against EGFP (Cas9), NESTIN and GFAP. CAS9 is widely expressed in the whole brain and co-localize with NESTIN and GFAP in the subventricular zone. *Left panels*: whole brain section; *right panels*: higher magnification of the left panel inset. Scale bars: left panels, 500 µm; right panels 100 µm. LV,lateral ventricle.

The Nestin-Cre is expressed quite early during development, beginning at E9.5, while the hGFAP-Cre appears to be expressed around E12.5-E13.5 ^10,11^. Both strains lead to widespread expression of the Cas9-P2A-EGFP throughout the brain of adult mice and pups (Fig. 1a-b and Supplementary Fig. 1b). Also of note it is the co-localization of NESTIN and GFAP with EGFP in the area of the sub-ventricular zone (Fig. 1a-b), one of the known site of neurogenesis of adult mice, indicating robust Cas9 expression in the NSC compartment.

### Efficient gene knockouts with RCAS-gRNA plasmids drive high-grade glioma formation

To explore whether the newly engineered RCAS/tv-a/Cas9 strains where suitable for *in vivo* genome editing, we first generated a series of RCAS plasmids that would allow the expression of gRNAs. For this purpose, we sub-cloned into the RCAS vector a cassette carrying a human U6 promoter (hU6), followed by a PGK promoter that drove the expression of a puromycin resistance gene (Puro) linked to a blue fluorescent protein (BFP) via a self-cleavable T2A peptide (hU6-gRNA-PGK-Puro-T2A-BFP) (Fig. 2a). We then cloned different previously described gRNAs targeting tumors suppressor genes (TSGs) frequently altered in high-grade gliomas: *Trp53*, *Cdkn2a* and *Pten*. All the RCAS plasmids generated for our studies were constructed using intermediate vectors compatible with the Gateway cloning system and the previously described RCAS-destination vector ^16^ (see Methods for details). To test the knockout efficiency of the RCAS-gRNA plasmids, we derived NSCs from *Ntv-a; LSL-Cas9* and infected them with a Cre-expressing plasmid to induce Cas9 expression. In parallel we also generated, by retroviral infection, NIH3T3 mouse fibroblasts expressing both TVA and the Cas9 genes. We then infected both cell lines with multiple rounds of infections using the various RCAS-gRNA plasmids. After either drug-selection (for the NSCs TVA-Cas9) or fluorescent activated cell sorting (FACS) (for the BFP in the NIH-3T3 TVA-Cas9) we verified the deletion of *Trp53*, *Cdkn2a* and *Pten* by western blot analysis. Since NIH-3T3 cells are *Cdkn2a* null, we tested the *Cdkn2a* gRNAs only in the NSCs. As shown in figure 2b, we observed efficient deletion of all those genes in both cellular systems.

**Figure 2:**
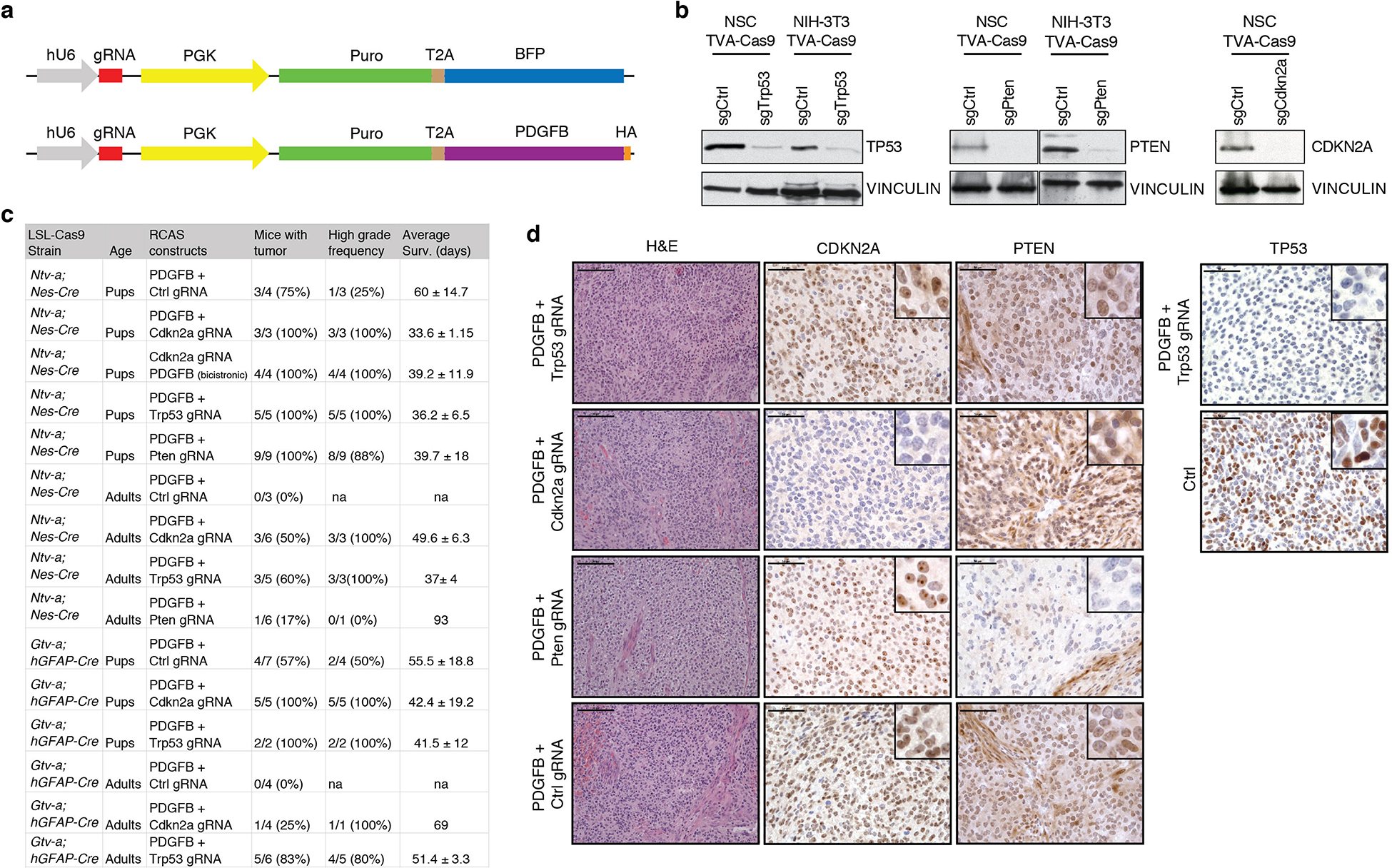
Tumor suppressor genes knockout by RCAS-gRNA plasmids induce high-grade-gliomas. **(a)** Schematic illustration of the RCAS-gRNA plasmids. **(b)** *In vitro* validation of the RCAS-gRNA against *Trp53*, *Pten, Cdkn2a* and a non-targeting control (Ctrl). Western blot analysis, using the indicated antibodies, on whole cell extracts from NIH3T3 TVA-Cas9 fibroblasts and *Ntv-a; LSL-Cas9* neural stem cells (NSCs) transduced with pMSCVhygro-CRE (NSC TVA-Cas9). To induce TP53 expression, the cells were collected 24h after exposure to ionizing radiation (10 Gy). **(c)** Table summarizing the injections performed in the *Ntv-a; Nes-Cre; LSL-Cas9* and *Gtv-a; hGFAP-Cre; LSL-Cas9* pups and adult mice. Co-injection of RCAS-PDGFB and the RCAS-gRNA against different tumor suppressor genes accelerate tumor formation, increases the tumor penetrance and the frequency of high-grade gliomas. **(d)** Hematoxylin and eosin (H&E) and immunohistochemical stainings (IHCs), using the indicated antibodies, of representative RCAS-PDGFB/gRNA tumors. To note PTEN expression in the normal vasculature but not in the tumor cells of the RCAS-PDGFB + RCAS-Pten-gRNA tumor. Insets show higher magnification images. Scale bars: H&E 100 µm; IHCs 50 µm.

We then tested the ability of the *Trp53*, *Cdkn2a* and *Pten* gRNAs to cooperate with PDGFB to induce high-grade gliomas (GBM) when injected into the *Ntv-a; Nes-Cre; LSL-Cas9* and *Gtv-a; hGFAP-Cre; LSL-Cas9* mice. RCAS-PDGFB intracranial injection into *Ntv-a* and *Gtv-a* pups has been previously shown to be sufficient to induce gliomas with variable penetrance (from 40% to 75%), but only a small fraction of the injected mice (25%) presented high-grade tumor features, such as pseudopalisades necrosis and microvascular proliferation ^17–19^. Moreover, RCAS-PDGFB injection into *Ntv-a* and *Gtv-a* adult mice resulted in very low tumor penetrance (approximately 15-20%) and long latency (over 100 days) ^13^. Co-injections RCAS-PDGFB and RCAS-TSG-gRNA (either one of *Trp53*, *Cdkn2a* or *Pten* gRNAs) into *Ntv-a; Nes-Cre; LSL-Cas9* and *Gtv-a; hGFAP-Cre; LSL-Cas9* pups resulted in a shortened tumor latency and increased total tumor incidence as compared to the co-injections of RCAS-PDGFB and RCAS-gRNA non-targeting control (Ctrl) (Fig. 2c and Supplementary Fig. 2). Most importantly, the vast majority (80-100%) of the RCAS-PDGFB/RCAS-TSG-gRNA injected mice showed histological features of high-grade gliomas (Fig. 2c-d). We also generated RCAS plasmids expressing the gRNA and the PDGFB in the same constructs (hU6-gRNA-PGK-Puro-T2A-PDGFB) (Fig. 2a). When injected into *Ntv-a; Nes-Cre; LSL-Cas9* pups, the RCAS-Cdkn2a-gRNA-PDGFB bicistronic vector was able to induce high-grade tumor formation with full penetrance and very short latency (approximately 40 days) (Fig. 2c and Supplementary Fig. 2).

Analogously to what was previously reported for the RCAS-PDGFB, injection in adult mice showed a considerably reduced tumor incidence. Actually, in our 120 days’ experimental timeframe, we didn’t observe any tumors in the mice co-injected with the RCAS-PDGFB and RCAS-gRNA non-targeting control neither in the *Ntv-a; Nes-Cre; LSL-Cas9* nor in the *Gtv-a; hGFAP-Cre; LSL-Cas9*. However, similarly to the injections in the pups, the injection of RCAS-PDGFB/RCAS-TSG-gRNA in adult mice lead to increased tumor incidence, with the majority of the tumor presenting high-grade characteristics (Fig. 2c and Supplementary Fig. 2).

Immunohistochemical (IHC) analysis of paraffin-embedded tissue showed loss of Trp53, Cdkn2a and Pten expression in the tumors injected with the corresponding RCAS-TSG-gRNA plasmid (Fig. 2d).

In summary, these data demonstrate that RCAS-gRNA constructs can induce deletion of the gene of interest in an *in vivo* setting and they could be used to efficiently target virtually any tumor suppressor gene.

### Inducible Cas9 expression in adult mice does not activate a robust immune response

The injection of the RCAS-TSG-gRNA plasmids into *Ntv-a; Nes-Cre; LSL-Cas9* and *Gtv-a; hGFAP-Cre; LSL-Cas9* mice should lead to an early deletion of the tumor suppressor gene of interest, due to the Cas9 expression at the time of the injection. Therefore, in order to have a mouse model in which we could generate the deletion of a specific gene in a time-controlled manner, to investigate for example the role of a gene not only in tumor initiation but also in tumor progression, we crossed the *Ntv-a; LSL-Cas9* mice with the *hUBC-CreERT2*, for inducible Cas9 expression upon tamoxifen exposure ^12^.

There have been controversial reports of immune response to Cas9 in some experimental models. On one hand, sign of both humoral and cellular immunity against Cas9 following systemic Adenoviral (Ad) vector-mediated Cas9 delivery was detected in mice ^20^. Moreover, injection of adeno-associated viruses (AAV) expressing Cas9 in the adult tibialis anterior muscle produced the elevation of CD45^+^ hematopoietic cells in the injected muscle and enlargement of the draining lymph nodes ^21^. On the other hand, Cas9 ribonucleoparticles (RNP) injection into the brain of adult mice showed an undetectable to mild microglia-mediated innate immune response ^22^.

As previously discussed above, Nestin-Cre and hGFAP-Cre are expressed quite early during embryogenesis, thus Cas9 expression under the control of those promoters is not expected to induce any immune response. However, we could not exclude such response upon tamoxifen-induced Cas9 expression in adult *Ntv-a; LSL-Cas9*; *hUBC-CreERT2* mice. Therefore, we performed an in-depth analysis of a possible immune response in the *Ntv-a; LSL-Cas9*; *hUBC-CreERT2* upon Cas9 induction. For this purpose, 4 weeks old *Ntv-a; LSL-Cas9*; *hUBC-CreERT2*^*+/+*^ and *Ntv-a; LSL-Cas9*; *hUBC-CreERT2*^*+/T*^ mice were treated with tamoxifen-containing diet for a total of 5 weeks. Whole blood samples were taken every two weeks. At the end of the experiment (week 9), blood, spleen and brain tissue were harvested and analyzed by flow cytometry, real-time quantitative PCR (qPCR) and immunofluorescence (Supplementary Fig. 3a). After 5 weeks of tamoxifen treatment we observed high percentage of EGFP positive cells in the blood (around 50%) and lower number in spleen and brain (approximately 10-15%) (Fig. 3a-b Supplementary Fig. 3c). Mice did not show any signs of inflammation and splenomegaly was not observed. Circulating levels of T-cells, B-cells, granulocytes and monocytes were determined by flow cytometry in the blood and spleen. While there were no significant differences neither in T cells (CD3^+^CD4^+^) nor B cells (CD3^-^B220^+^), there was a trend towards decreased circulating Gr-1 positive neutrophils in the blood of the *Ntv-a; LSL-Cas9*; *hUBC-CreERT2*^*+/T*^ tamoxifen-treated mice, which could signify inefficient production of these cells in the bone marrow (Fig. 3a). Despite this reduction, we did not detect significant differences in neither the number nor the percentage of granulocytes in whole blood cell counts (Supplementary Fig. 3b). Flow cytometry in the brain showed no changes in lymphocytes, microglia or macrophages with the gating strategy described previously ^23^ using CD45 and CD11b. We further confirmed by qPCR that there were no major differences in mRNA expression of a panel of microglia activation specific markers (*CD45*, *IL12a-1*, *P2ry12*, *Tmem119*, *Cx3cr1* and *Iba-1*) (Supplementary Fig. 3d).

**Figure 3:**
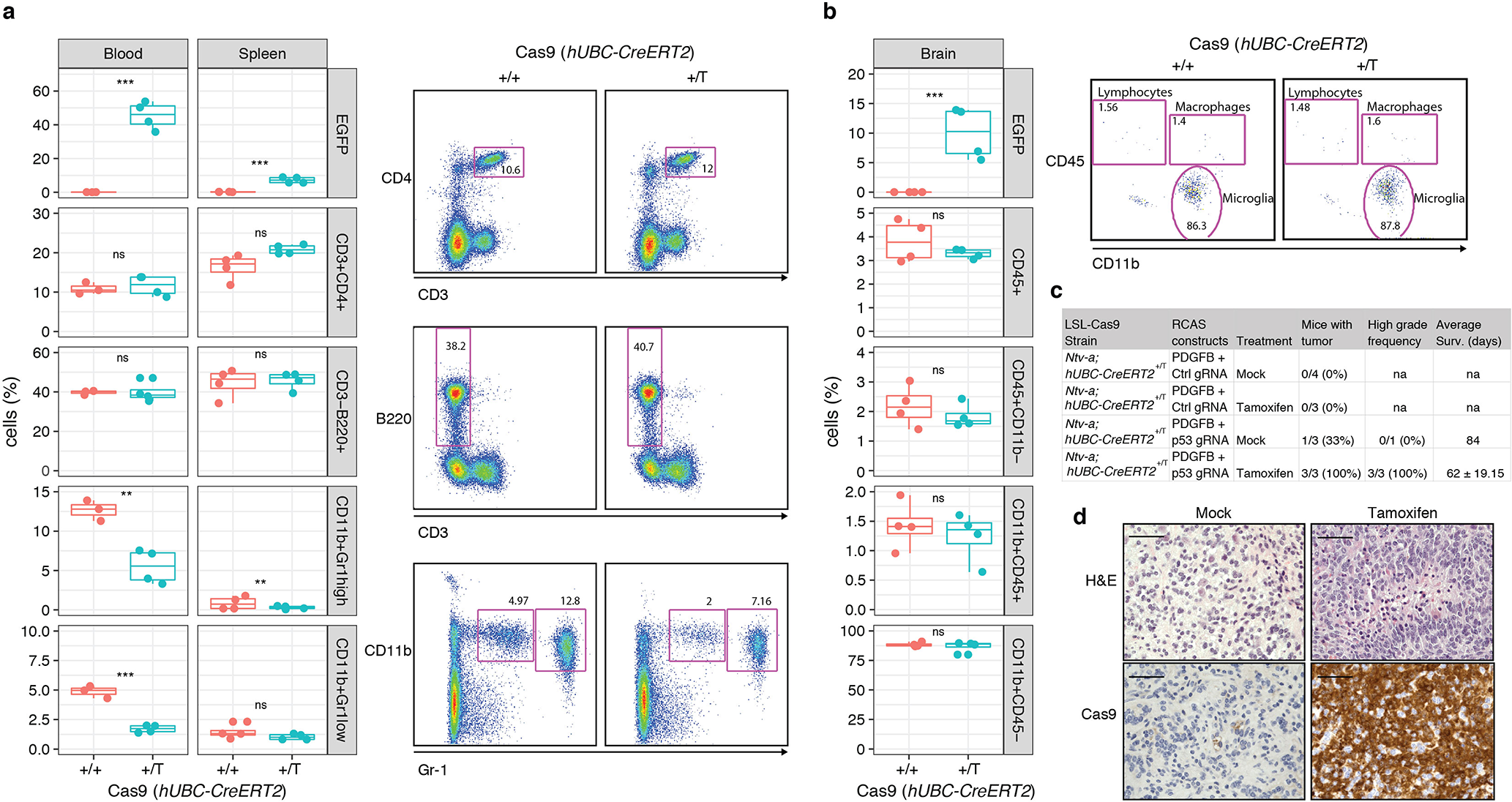
Ubiquitous Cas9 expression in TVA/Cas9 adult mice does not induce a robust immune response. **(a)** *Left panels*: Flow cytometry analysis for the specified markers in blood and spleen of *Ntv-a; LSL-Cas9*; *hUBC-CreERT2* mice of the indicated genotype. Four weeks old mice were treated with tamoxifen in the food for 5 consecutive weeks. *Right panels*: representative flow cytometry plots, with the gating strategy used for the analysis. **(b)** *Left panels*: Flow cytometry analysis for lymphocytes, macrophages and microglia of the brain of the mice in (a). *Right panels*: representative flow cytometry plots, with the gating strategy used for the analysis. **(c)** Table summarizing the injections performed in the *Ntv-a; LSL-Cas9*; *hUBC-CreERT2* adult mice. Mice injected with RCAS-PDGFB + RCAS-Trp53-gRNA and treated with tamoxifen to induce Cas9 expression, develop high-grade gliomas. **(d)** H&E and Cas9 IHCs of RCAS-PDGFB + RCAS-Trp53-gRNA tumors for the indicated treatment. High-grade glioma features and CAS9 expression are present only in the tamoxifen-treated mice. Scale bars: H&E 100 µm; IHCs 50 µm.

Since our analysis did not suggest an immune response to Cas9 expression in the *Ntv-a; LSL-Cas9*; *hUBC-CreERT2*^*+/T*^ mice, we proceeded to inject them with the RCAS-PDGFB/RCAS-gRNA. Four weeks old adult mice were injected intracranially with the RCAS-PDGFB in combination with either RCAS-p53-gRNA or RCAS-Ctrl-gRNA. Two weeks after injection, the mice were separated in two groups and treated for two weeks with either mock-treatment or tamoxifen (see Methods for details). Mice were then sacrificed either upon sign of tumor development or at the end of the experiment (90 days). As for the *Ntv-a; Nes-Cre; LSL-Cas9* and *Gtv-a; hGFAP-Cre; LSL-Cas9* strains, none of the *Ntv-a; LSL-Cas9*; *hUBC-CreERT2*^*+/T*^ mice injected with the RCAS-PDGFB and RCAS-Ctrl-gRNA developed tumors (Fig. 3c). While only one out of 3 of the mock-treated *Ntv-a; LSL-Cas9*; *hUBC-CreERT2*^*+/T*^ mice injected with the RCAS-PDGFB and RCAS-p53-gRNA developed a low-grade tumor at 84 days, all the mice treated with tamoxifen were sacrificed at earlier time due to high-grade gliomas (Fig. 3c-d).

### The *Bcan-Ntrk1* gene fusion produce high-grade gliomas

Gene fusions have been documented as cancer-drivers for more than three decades, providing valuable insights into the tumorigenesis process. The occurrence and importance of gene fusions in glioma has been appreciated only recently, largely due to high-throughput technologies, and gene fusions have been indicated as one of the major genomic abnormalities in GBM ^24^. The functional role of the vast majority of these alterations is completely unexplored. Recurrent gene fusions involving the Trk receptor family (NTRK1, 2 and 3 genes) have been recently described in a variety of tumors, including gliomas ^25–27^. Here, we decided to focus on the *BCAN-NTRK1* gene fusion, identified in glioblastoma and glioneuronal tumors ^28,29^.

*BCAN* and *NTRK1* are located on chromosome (Chr) 1 q23.1 and the *BCAN*-*NTRK1* fusion gene results from of an intra-chromosomal deletion that juxtapose the *BCAN* exon 13 with the *NTRK1* exon 11 (Supplementary Fig. 4a). The *BCAN* gene codes for Brevican, a glycoprotein that is highly expressed in the brain, while *NTRK1*, that codes for the TrkA kinase, is almost undetectable in the adult brain (Supplementary Fig. 4b). The mouse homologues, *Bcan* and *Ntrk1*, located on Chr3, have a similar gene structure and expression pattern to their human counterparts (Supplementary Fig. 4a-b). Hence, we argued that the *Bcan*-*Ntrk1* fusion would be an appropriate genomic alteration to be studied with the RCAS/tv-a/Cas9 system.

**Figure 4:**
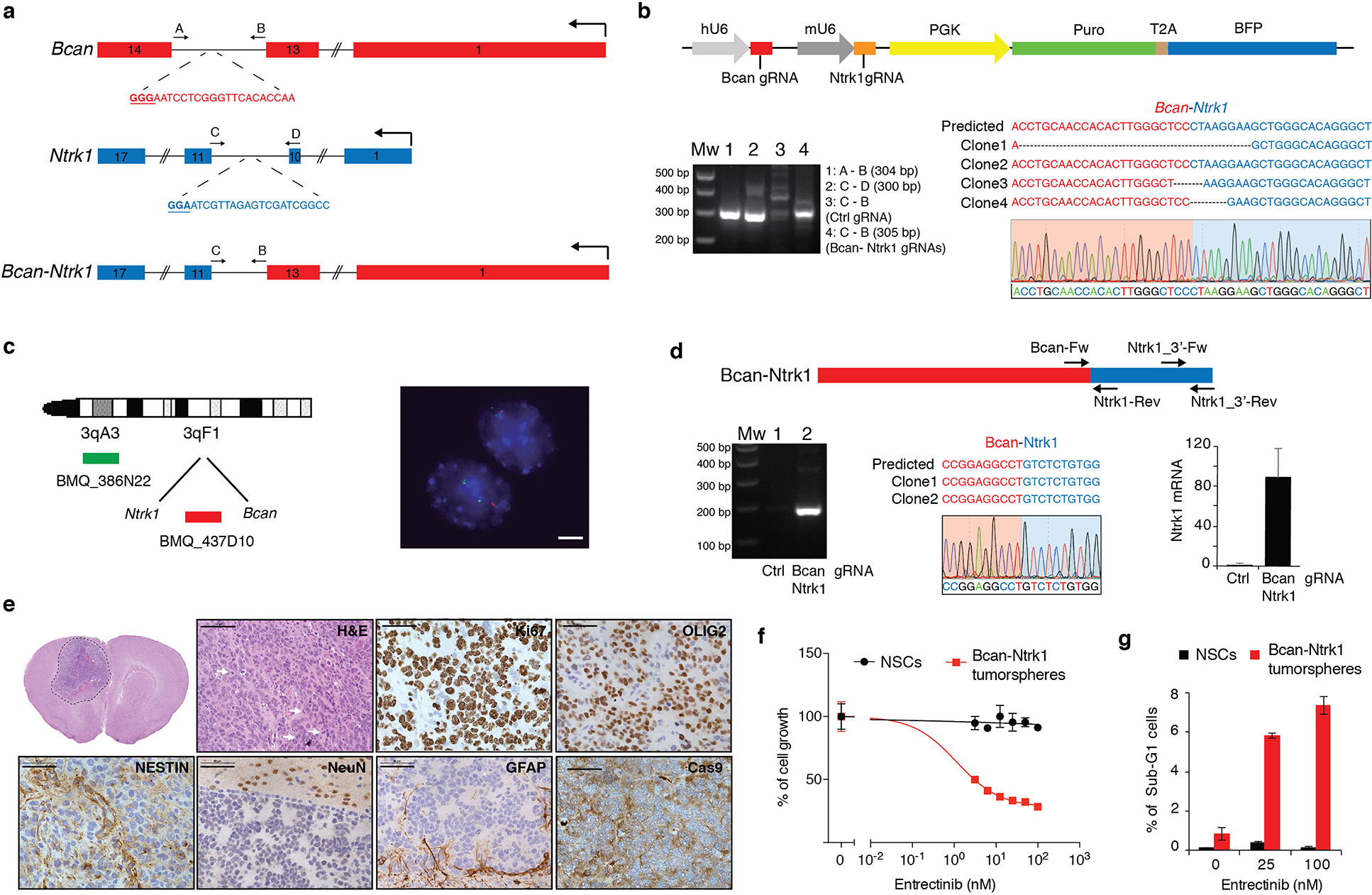
*Bcan-Ntrk1* gene fusion drives high-grade glioma formation. **(a)** Schematic representation of the *Bcan* and *Ntrk1* gene loci and the *Bcan-Ntrk1* gene fusion. Indicated are the gRNAs targeting both genes and the primers used for the PCR amplification of the indicated genomic regions. **(b)** *Top panel*: RCAS-gRNA-pair vector expressing the Bcan and Ntrk1 gRNAs. *Bottom panels*: PCRs were performed with the specified primers on genomic DNA extracted from the *p53-null* TVA-Cas9 NSCs transduced with the indicated gRNAs. The PCR band for the Bcan-Ntrk1gRNA infected cells was sub-cloned and analyzed by Sanger sequencing. The sequences of four independent clones and a representative chromatogram are shown. **(c)** *Left panel*: Diagram of fluorescence *in situ* hybridization (FISH) probe design. BAC clone BMQ-437D10 (red) is located within the deleted region and BMQ-386N22 (green) is used as a control of chromosome 3. Mouse BACs are represented as green and red bars. *Right panel:* Representative FISH results using the two-color probe designed to detect the NtrK1-Bcan intergenic microdeletion. The control green signal was used to count the number of chromosomes 3. The loss of the red signals indicates the microdeletions. Scale bar: 5 µm. **(d)** *Top panel*: Schematic representation of *Bcan-Ntrk1* fusion transcript. *Bottom panels*: (*left*) RT-PCRs were performed on the mRNA from the *p53-null* TVA-Cas9 NSCs transduced with the indicated gRNAs, using the Bcan-Fw and Ntrk1-Rev primers; (*middle*) the PCR band was sub-cloned and the sequences of 2 independent clones and a representative chromatogram are shown; (*right*) quantitative real-time PCR (qPCR), with the Ntrk1_3’-Fw and Ntrk1_3’-Rev primers, showing the upregulation of the Ntrk1 mRNA in the cells expressing the *Bcan-Ntrk1* fusion. **(e)** H&E and IHCs using the indicated antibodies. White arrows point to mitotic figures. Scale bars: H&E 100 µm; IHCs 50 µm. **(f)** Cell proliferation assay performed on *p53-null* TVA-Cas9 NSCs and Bcan-Ntrk1 tumorspheres exposed for 96h to increasing doses of Entrectinib, a pan-Trk inhibitor. **(g)** Entrectinib treatment in Bcan-Ntrk1 tumorspheres induces an increase in apoptosis, as measured by percentage of sub-G1 population in cell fixed and stained with propidium iodide.

In order to generate the *Bcan*-*Ntrk1* gene fusion we designed gRNAs in the introns 13 and 10 of *Bcan* and *Ntrk1,* respectively (Fig. 4a). The pair of gRNAs was subsequently cloned into an RCAS plasmid containing both a hU6 and mU6 promoters (hU6-gRNA-mU6-gRNA-PGK-Puro-T2A-BFP) (RCAS-gRNA-pair) (Fig. 4b, *top panel*), with a previously described strategy ^30^. The RCAS-gRNA-pair vector was then used to infect the NIH-3T3 TVA-Cas9 (data not shown) and also *p53-null* TVA-Cas9 NSCs isolated from *Gtv-a; hGFAP-Cre; LSL-Cas9; p53*^*lox/lox*^ pups. Generation of the expected chromosomal deletion was tested by PCR on genomic DNA, and later analyzed by sequencing (Fig. 4b, *bottom panel*). Furthermore, we used fluorescence *in situ* hybridization (FISH) to evaluate the frequency of cells carrying the desired deletion on Chr3. Approximately 40% of the cells (80/209) showed the loss of one copy of the probe located between the *Ntrk1* and *Bcan* gene, indicating that the generation of the *Bcan*-*Ntrk1* rearrangement is a relatively efficient process (Fig 4c).

We then confirmed the expression of the *Bcan*-*Ntrk1* fusion transcript, in the *p53-null* TVA-Cas9 NSCs, by reverse transcription PCR and sequencing of a cDNA fragment overlapping the fusion exon junction (Fig. 4d). Analogously to what has been observed in the GBM patients carrying the *BCAN*-*NTRK1* fusion (Supplementary Fig. 4c), the generation of the *Bcan*-*Ntrk1* rearrangement leads to exceptionally high levels of the *Ntrk1* 3’ mRNA region involved in the gene fusion (Fig. 4d, *bottom right panel*).

To test whether the *Bcan*-*Ntrk1* gene fusion was sufficient to drive glioma formation we injected intracranially into *NOD/SCID* mice the *p53-null* TVA-Cas9 NSCs infected with the RCAS-gRNA-pair. While none (0/5) of the mice injected with the control NSCs developed tumors during the observation period (90 days), 4 out of the 6 mice injected with the *Bcan*-*Ntrk1*gRNA pairs had to be sacrificed due to sign of tumor formation (with mean survival of 72 ± 14 days). Histopathological examination of the tumors evidenced a series of characteristics typical of high-grade gliomas: nuclear atypia, high number of mitotic figures, necrotic areas and infiltration in the normal brain parenchyma (Fig. 4e and Supplementary Fig. 4d-e). *Bcan*-*Ntrk1-* induced tumors showed elevated percentage of Ki67 positive cells, were positive for OLIG2 and NESTIN, negative for the neuronal marker NeuN and, besides the small percentage of astrocytes trapped inside the tumor, GFAP positive cells were almost exclusively detected at the normal/tumor border (Fig. 4d). Moreover, IHC staining evidenced high level of expression of Ntrk1 as compared to a PDGFB-induced tumor (Supplementary Fig. 4f).

We further confirmed by genomic PCR and FISH analysis the presence of the *Bcan*-*Ntrk1* gene fusion on cells isolated from the tumor-bearing mice, propagated *in vitro* as tumorspheres (Supplementary Fig. 4g-h). Strikingly, these tumorspheres expressed very high levels of *Ntrk1* as compared to both the NSCs control or to the *Bcan*-*Ntrk1* NSCs prior intracranial injection (Supplementary Fig. 4i). These data would suggest that *in vivo,* from the mixed population of the NSCs infected with the *Bcan*-*Ntrk1* gRNAs, those cells that carried the gene rearrangement were positively selected.

There has been a lot of interest lately in targeting *NTRK* gene fusions across multiple tumor types ^31^. Entrectinib is a first-in-class pan-TRK kinase inhibitor currently undergoing clinical trials in a variety of cancers. To confirm that the *Bcan*-*Ntrk1* tumors we generated were dependent on TrkA activity, we treated *in vitro* with Entrectinib the *Bcan*-*Ntrk1* tumorspheres. As shown in figure 4f, the *Bcan*-*Ntrk1* tumorspheres were exquisitely sensitive to Trk inhibition, while no effect was observed on the control *p53-null* TVA-Cas9 NSCs. Entrectinib led to a significant reduction of tumor cells growth associated with an increase of the number of apoptotic cells, detected as sub-G1 population in a propidium iodide staining (Fig. 4f-g).

Overall these data indicate that the RCAS/TVA-CRISPR/Cas9 system is a very powerful model to study the role of gene fusions in tumorigenesis and as possible therapeutic targets.

### Generation of the *Myb-Qk* chromosomal translocation

The human *BCAN-NTRK1* and mouse *Bcan*-*Ntrk1* fusions are generated by a small chromosomal deletion of approximately 200Kb. To test whether the RCAS/tv-a/Cas9 system was also suited for inter-chromosomal translocations, we decided to model the *MYB-QKI* gene fusion, a recently identified putative driver of a subtype of pediatric low-grade gliomas (PLGG), known as angiocentric gliomas ^32^. Although *MYB* and *QKI* are both located on Chr6 in human, the mouse homologues *Myb* and *Qk* are located on different chromosomes, Chr10 and Chr17, respectively.

*MYB* encodes for a transcription factor that is a key regulator of hematopoietic cell proliferation and deregulated MYB activity has been observed in variety of human cancers. *QKI* is a tumor suppressor gene that encodes for a RNA-binding protein, QUAKING, that plays a role in the development of the CNS, among other organs. Several *MYB-QKI* gene fusions have been described in angiocentric gliomas, all of them involved the same *QKI* 3’ region (exon 5 to 8) fused to different MYB exons (1-9, 1-11 or 1-15). Here, we focused on the most frequent *MYB* (exon1-9) *-QKI* (exon 5 to 8) fusion event.

To generate the mouse *Myb* (exon1-9) *- Qk* (exon 5 to 8) fusion, we designed gRNAs in the intron 4 for *Myb* and 9 for *Qk* (Fig. 5a), and we cloned them into the RCAS-gRNA-pair. Genomic PCR and sequencing from the NIH-3T3 TVA-Cas9 (data not shown) and also from *p53-null* TVA-Cas9 NSCs infected with the RCAS-Qk-gRNA-Myb-gRNA, confirmed the generation of the *Myb-Qk* fusion (Fig. 5b). By RT-PCR we also observed the expression of the *Myb-Qk* transcript (Fig. 5c).

**Figure 5:**
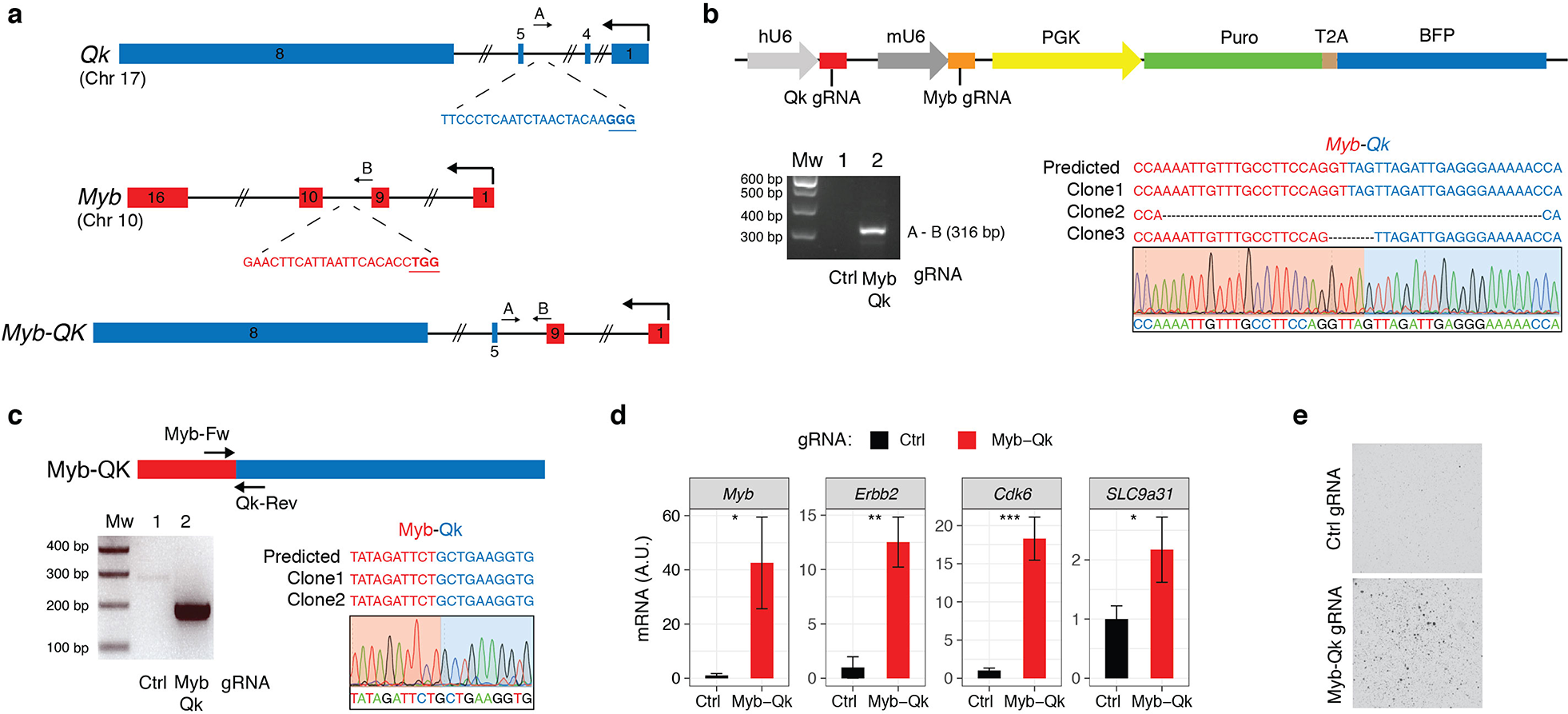
Generation of the *Myb-Qk* chromosomal translocation. **(a)** Schematic representation of the *Myb* and *Qk* gene loci and the *Myb-Qk* gene fusion. Indicated are the gRNAs targeting both genes and the primers used for the PCR amplification of the indicated genomic regions. **(b)** *Top panel*: RCAS-gRNA-pair vector expressing the Myb and Qk gRNAs. *Bottom panels*: PCRs were performed with the specified primers on genomic DNA extracted from the *p53-null* TVA-Cas9 NSCs transduced with the indicated gRNAs. The PCR band for the Myb-Qk gRNA infected cells was sub-cloned and analyzed by Sanger sequencing. The sequences of three independent clones and a representative chromatogram are shown. **(c)** *Top panel*: Schematic representation of *Myb-Qk* fusion transcript. *Bottom panels*: (*left*) RT-PCRs were performed on the mRNA from the *p53-null* TVA-Cas9 NSCs transduced with the indicated gRNAs, using the Myb-Fw and Qk-Rev primers; (*right*) the PCR band was sub-cloned and the sequences of 2 independent clones and a representative chromatogram are shown. **(d)** qPCR analysis shows the upregulation of Myb-activated genes in the *p53-null* TVA-Cas9 NSCs expressing the *Myb-Qk* fusion. **(e)** *p53-null* TVA-Cas9 NSCs transduced with the Myb and Qk gRNAs, but not Ctrl gRNA, are able to growth in soft agar.

In human and mouse normal adult brain, Myb mRNA expression is almost undetectable (Supplementary Fig. 5b). The *MYB-QKI* fusion has been shown to functionally activate the *MYB* promoter and to possibly contribute to an autoregulatory feedback loop ^32^. Indeed, when we measured *Myb* expression in cells expressing the *Myb-Qk* fusion we observed an increase of Myb mRNA as compared to control cells (Fig. 5d). We also observed increased mRNA levels of a series of genes (*Erbb2*, *Cdk6* and *Slc9a31*) that have been shown to be upregulated by the *MYB-QKI* fusion^32^ (Fig. 5d).

We then tested their transforming potential *in vitro* by plating the cells in soft-agar, and we observed that the *p53-null* NSCs expressing the *Myb-Qk* fusion, but not the *p53-null* NSCs infected with the Ctrl gRNA cells, were able to form colonies.

### Modeling BRAF V600E mutation by homology directed repair (HDR)

One of the known applications of the CRISPR/Cas9 system is to induce point mutations through Homologous Recombination (HR). Delivery of a gRNA with either double-stranded DNA (dsDNA) or single-stranded DNA (ssDNA) repair templates, containing a desired modified sequence together with variable length upstream and downstream homology arms, has been used to recreate oncogenic driver mutations ^4^.

Activating mutations in the *BRAF* kinase gene (V600E) have been identified in various types of pediatric gliomas (Pilocytic astrocytomas (< 10%), pleomorphic xanthoastrocytomas (WHO grades II and III; 50%–65% cases), gangliogliomas (20%–75% cases)) and also adult high-grade gliomas (5%) ^33^.

To model a missense gain-of-function *Braf* mutation we used the strategy previously described to generate a *Kras*^*G12D*^ mutation ^4^ and designed an HDR donor template, which comprises of an 800bp genomic sequence covering exon 18 of the mouse *Braf* gene. This HDR donor encoded: (i) a valin (V) to glutamine (E) mutation in the amino acid position 637 (V637E), resulting in the oncogenic *Braf*^*V637E*^ mutation, homologous to the human *BRAF*^*V600E*^; (ii) 11 synonymous single-nucleotide changes to discriminate the difference between the donor and wild-type sequences and to mutate the protospacer-adjacent motif (PAM) to avoid donor DNA cleavage by Cas9. The HDR donor template, together with a gRNA targeting a sequence 22bp upstream the Braf V637 residue, were subsequently cloned into a lentiviral vector (Fig. 6a) and transduced into the *p53-null* TVA-Cas9 NSCs. We then evaluated the efficiency of HDR-mediated *Braf* mutation by PCR, sub-cloning of the amplified cDNA region and Sanger sequencing. Sixty percent (6/10) of the analyzed clones contained the desired V637E mutation and also a second point mutation D624N (Supplementary Fig. 6a). This latter mutation, although undesired, is a conservative mutation from an aspartate to an asparagine residue and it’s not expected to have any functional consequence on BRAF activity.

**Figure 6:**
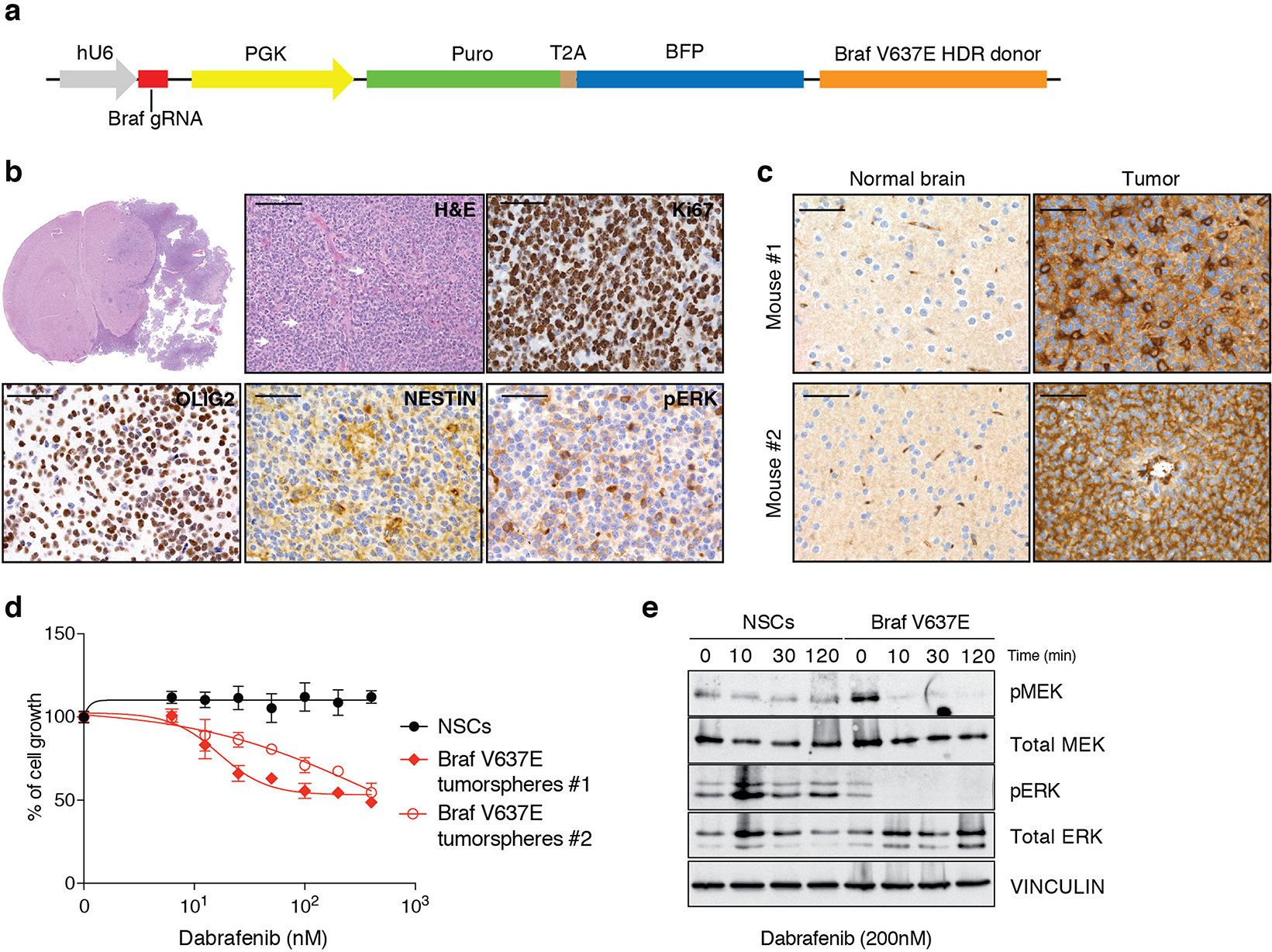
Glioma formation induced by *Braf* ^*V637E*^. **(a)** Schematic representation of the plasmid carrying the Braf gRNA and the Braf^V637E^ Homolgy-Directed-Repair (HDR) donor. **(b)** H&E and IHCs using the indicated antibodies. White arrows point to mitotic figures. Scale bars: H&E 100 µm; IHCs 50 µm. **(c)** IHCs using an anti-BRAF V600E antibody on two Braf^V637E^ mutant tumors. Contralateral normal brain was used as a negative control. **(d)** Cell proliferation assay performed on *p53-null* TVA-Cas9 NSCs and *Braf*^*V637E*^ tumorspheres exposed for 96h to increasing doses of Dabrafenib. **(e)** Western blot analysis using the specified antibodies on *p53-null* TVA-Cas9 NSCs and *Braf*^*V637E*^ tumorspheres grown for 24h in absence of growth factors and then treated with Dabrafenib (200nM) for the indicated time.

When transplanted intracranially into *NOD/SCID* mice, the *p53-null Braf*^*V637E*^ NSCs induce tumor formation in 100% of the injected mice (6/6), with an average survival of 66 ± 11.5 days. Histopathological examination of the tumors evidenced a number of features characteristic of high-grade gliomas: nuclear atypia, high number of mitotic figures and necrotic areas (Fig. 6b and Supplementary Fig. 6b). Moreover, we observed clusters of tumor cells infiltrating the normal brain parenchyma, with the vast majority of these cells surrounding tumor vessels (Supplementary Fig. 6b), resembling the vascular co-option observed both in primary and metastatic brain tumors. It is also to note the presence of some giant cells (Supplementary Fig. 6c) and areas of the tumors with epithelioid morphology (Supplementary Fig. 6d).

Immunohistochemical analysis of the BRAF mutant tumors revealed high percentage of Ki67 positive cells, positivity for both OLIG2 and NESTIN and elevated MAPK kinase activity, as evidenced by pERK IHC (Fig. 6b). Additionally, we were able to confirm the expression of the BRAF V637E mutation using an antibody specifically designed to recognize the human BRAF V600E mutant.

To validate that the BRAF mutant tumors were dependent on an active BRAF signaling pathway, we isolated tumorspheres from two of those tumors and we treated them *in vitro* with Dabrafenib, a specific BRAF inhibitor that is currently in clinical trials for *BRAF*^*V600E*^ mutant melanomas. Both tumorspheres lines carried the *Braf*^*V637E*^ mutation as confirmed by PCR and Sanger sequencing (Supplementary Fig. 6e). As shown in figure 6d, Dabrafenib treatment induced growth reduction in both *Braf*^*V637E*^ tumors, but not in *p53-null* NSCs. Moreover, western blot analysis showed a reduction of MAPK kinase signaling pathway after exposure to Dabrafenib in *Braf*^*V637E*^ tumor cells but not in control cells (Fig. 6e).

In summary, we have presented a platform that will be useful to study not only the role of tumor suppressor genes and genomic rearrangements but also of potential oncogenic mutations.

## Discussion

Here we have established a novel powerful methodology for precision tumor modeling *in vivo* and *ex vivo*, by combining the versatility of the genome editing CRISPR/Cas9 technology with the specificity of somatic gene transfer mediated by the RCAS/TVA system.

The latest improvements in the genetically engineered mouse modeling (GEMM) have contributed to the understanding of the molecular pathways responsible for tumor initiation and progression. Few elements should be taken in consideration to properly mimic the natural history of a tumor: a) introduction of the same mutations found in human tumors, ideally in their endogenous loci; b) the genetic alterations should be silent during embryonic and early postnatal development (with the exception of models of familiar or pediatric tumors); c) the mutant genes should be expressed in particular target tissues or in specific cell types and e) the mutations should be present in a limited number of cells. The RCAS/TVA-CRISPR/Cas9 system described here gives the possibility to recapitulate all of these characteristics in one single model.

A number of different knockin TVA mouse models have been published in the past and they have been used to study a variety of cancers: gliomas, medulloblastomas, melanoma, breast, pancreatic, ovarian and liver cancer ^7,34^. Breeding of any of the TVA lines to the knockin Cas9 strain would allow to generate novel somatic genome editing models to study a plethora of tumor types. Moreover, Seidler and colleagues have recently described a Cre-dependent TVA-transgenic line (*LSL-R26-*^*TVA-lacZ*^) ^35^ that in combination with the Cas9 knockin mice and one of the hundreds cell-type-specific Cre mouse strains ^36,37^ would allow the use of the RCAS/TVA-CRISPR/Cas9 system for somatic genome editing virtually in any proliferative cells of the organism.

As a proof-of-principle, we used two different CNS-specific TVA-transgenic lines (*Nestin-tv-a* and *GFAP-tv-a*) to perform functional characterization of various genetic alterations described in human gliomas: 1) knockout of a panel of TSGs recurrently lost or mutated in GBMs (*TP53*, *CDKN2A* and *PTEN*), 2) genomic rearrangements identified in different subtypes of gliomas (*BCAN*-*NTRK1* and *MYB*-*QKI*) and 3) a point mutation (BRAF^*V600E*^) present in a variety of pediatric and adult gliomas.

For the TSGs we selected *Trp53*, *Cdkn2a* and *Pten*, since they were previously shown to cooperate with PDGFB overexpression to induce high-grade gliomas ^13,18^. Indeed, co-injection of gRNAs targeting those genes led to the formation of GBMs with high frequency. These data would suggest that combining the expression of specific oncogene drivers with gRNAs for a TSG of interest would quickly provide information on its contribution to the tumorigenesis process. Moreover, by using the *hUBC-CreERT2*^*+/T*^ or other Cre-inducible strains it will be possible to exploit the RCAS/TVA-CRISPR/Cas9 system not only to study tumor initiation, but also tumor progression and maintenance.

Due to the quite recent advancement in the CRISPR/Cas9 technology, very few mouse models have been previously developed to study brain tumorigenesis ^38–40^. By *in utero* electroporation (IUE) of the forebrain of mouse embryos using plasmids encoding Cas9 in combination with gRNAs targeting *Nf1*, *Trp53* and *Pten*, Zuckerman and Chen were able to induce highly aggressive tumors that had histopatological features of human GBMs. More recently, Cook and colleagues have used adenoviral (Ad) vectors to express Cas9 and to generate the *BCAN-NTRK1* rearrangement in the brain of adult mice ^40^.

In our opinion there are at least two key issues with the use of IUE and Ad for glioma CRISPR/Cas9 modeling: timing of the gRNA delivery and lack of specificity of the targeted cells. Electroporation is normally performed at E14.5 or E15.5 and genetic alterations at this gestational stage might not be necessarily reflecting the biology of gliomas in the adult. The second issue is that the expression of the Cas9 enzyme from a constitutive promoter, as it has been used in both IUE and Ad studies, does not allow genome editing in a cell-type-specific manner hence not restricting the genetic alteration to the putative cells of origin of gliomas. This latter point is particularly relevant for proper cancer modeling, since it has been shown that the same driver mutations can lead to phenotypically and molecularly diverse glioma subtypes from different pools of adult CNS progenitor cells ^41,42^.

The CRISPR/Cas9 system has been previously used to model genomic rearrangements both in human and mouse cells ^43–45^. Here we generated the *Bcan*-*Ntrk1* gene fusion, via a microdeletion of 0.2Mb on Chr3, and the *Myb*-*Qk* gene fusion, by a chromosomal translocation. Fusion transcripts can generally lead to at least four different situations: a) increased overexpression of an oncogene (e.g. *IgH-MYC* in leukemia), b) deregulation of a tumor suppressor gene (e.g. *CHEK2-PP2R2A* in childhood teratoma), c) generation of a new aberrant protein (e.g. *BCR-ABL1* in leukemia), and d) a combination of various of the above (e.g. *MYB-QKI* in angiocentric glioma) ^32,46^.

We have observed that the *NTRK1* gene fusions lead to overexpression of the chimeric *NTRK1* transcripts in human glioma patients and in our mouse model (Supplementary Fig. 4c and 4i). Most likely, the very pronounced levels of TrkA kinase activity achieved by the high levels of the chimeric *NTRK1* transcripts are responsible for the oncogenic activity of those fusions. Indeed, we observed that the *Bcan*-*Ntrk1* tumors were finely sensitive to the pan-Trk inhibitor Entrectinib.

The *MYB-QKI* rearrangement has been shown to drive tumorigenesis through a tripartite mechanism: MYB activation by truncation, aberrant MYB-QKI expression and hemizygous loss of the tumor suppressor *QKI* ^32^. Using the RCAS/TVA-CRISPR/Cas9 system we successfully generated the *Myb*-*Qk* gene fusion in mouse cells and indeed we observed an increased *Myb* activation, as shown by upregulation of some Myb-regulated genes (*Erbb2*, *Cdk6* and *Slc9a31*) (Fig. 5d). Although it is conceivable that loss of the *Qk* gene is contributing to the tumorigenic potential of the cells carrying the *Myb*-*Qk* gene fusion, further work will be needed to clarify it.

Despite that the generation of point mutations with the CRISPR/Cas9 system might represent one of the most powerful feature of this genome editing technology, it is also the most challenging and very few cancer models have been developed with it ^4,47–49^. Here we generated the first CRISPR/Cas9 model for the BRAF V600E mutation.

BRAF V600E mutation has been identified in approximately 60% of pleomorphic xanthoastrocytomas (PXAs), as well as in varying percentages of other types of gliomas. Very interestingly, the tumors that carried the *Braf*^*V637E*^ knockin mutation (homologous to the human *BRAF*^V600E^ mutant allele) resemble the epithelioid variant of glioblastoma. The tumor cells exhibit epithelioid features and discohesiveness reminiscent of this entity. Epithelioid GBMs (E-GBM) feature high rates of BRAF V600E mutation and are thought to arise from the malignant transformation of PXAs ^50,51^. Here we have shown that *Braf*^*V637E*^ tumors are quite sensitive to Dabrafenib treatment, suggesting that this inhibitor might represent a possible therapeutic approach for those glioma types.

In conclusion, we have developed an extremely powerful and versatile mouse model that combines the somatic genome transfer ability of the RCAS/TVA system with the CRISPR/Cas9 genome editing technology. We believe that such a flexible model will greatly expedite the generation of precise cancer models.

## Methods

### DNA constructs, Design and Cloning of guide RNAs

The pKLV-U6gRNA-PGKpuro2ABFP (Plasmid #50946) ^52^, lentiCas9-Blast (Plasmid #52962) ^53^ and pMSCVhygro-CRE (Plasmid #34565) ^54^ were obtained from Addgene. The retroviral RCAS Gateway Destination Vector (RCAS-Y-DV) ^16^ was kindly provided by Eric Holland. To sub-clone the gRNA into the RCAS-Y-DV vector we generated a pDONR-gRNA plasmid, by a multiple steps process. First, the region containing the hU6 promoter, *Bbs*I cloning sites, gRNA scaffold, PGK promoter, and selectable markers Puromycin and BFP (Blue Fluorescent protein) (hU6-gRNA-PGK-Puro-T2A-BFP) was amplified by PCR from the pKLV-U6gRNA-PGKpuro2ABFP plasmid using the Platinum Pfx Kit (Invitrogen, Cat. 11708-013) and the primers aTTB Fw and aTTB Rv (Supplementary Table1). The PCR-amplified product was transferred by site-specific recombination (Gateway BP Clonase, Invitrogen, Cat. 11789-020) into the pDONR221 Vector (Invitrogen, Cat. 12536017) following manufacturer’s instructions. Lastly, the *Bbs*I restriction site at position 437 was removed by site-directed mutagenesis (QuikChange Lightning Site-Directed Mutagenesis kit, Agilent, Cat. 210518) using the primers pDONR_BbsI_mut-Fw and Rv (Supplementary Table1). This step was necessary to remove, from the pDONR221, a *Bbs*I restriction site outside the gRNA cloning site.

To generate the pDONR-gRNA that expressed also the PDGFB-HA, we performed a PCR using the Platinum Pfx Kit and the primers pDONR-Fw and Rv for the backbone and PDGFB-Fw and Rv for the insert (Supplementary Table1). The two fragments were then assembled using the Gibson Assembly Master Mix (New England Biolabs, Cat. E2611L). To obtain the final plasmid, the *Bbs*I restriction site in the PDGFB sequence (pDONR-sgRNA-PDGFB position 2822) was removed by site-directed mutagenesis using the primers PDGFB_ BbsI_mut-Fw and Rv.

The pDONR-gRNA plasmids were recombined into the RCAS-Y-DV using the Gateway LR Clonase II Enzyme mix (Invitrogen, Cat. 11791100), following the manufacturer’s instructions. All the constructs were verified by Sanger-sequencing.

The gRNA sequences targeting *Cdkn2a*, *Pten* and *Tp53* were previously described ^38,55,56^. *Bcan*, *Ntrk1*, *Myb*, *Qk* and *Braf* gRNAs were designed using the Genetic Perturbation Platform web portal (http://portals.broadinstitute.org/gpp/public/analysis-tools/gRNA-design).

For cloning of single gRNAs, oligonucleotides containing the *Bbs*I site and the specific gRNA sequences were annealed, phosphorylated and ligated either into the pDONR-gRNA or the pKLV-U6gRNA(BbsI)-PGKpuro2ABFP previously digested with *Bbs*I. The cloning of the paired gRNA was done according to the protocol described by Vidigal and colleagues ^30^. Briefly, the oligonucleotides containing the different gRNA-pairs (Supplementary Table1) were amplified with Phusion High-Fidelity polymerase (New England Biolabs, M0530S) using primer F5 and R1 (Supplementary Table1). PCR products were gel-purified and ligated to *Bbs*I-digested pDonor_mU6 plasmid (kindly provided by A. Ventura) by using the Gibson Assembly Master Mix (New England Biolabs 174E2611S). The Gibson reaction was then digested with *Bbs*I at 37°C for 3 h. The linearized fragment containing the pair gRNA, the mU6 promoter and the gRNA scaffold was gel-purified and cloned into the pDONR-gRNA and then into the RCAS-Y-DV.

The Braf V637E HDR donor (Supplemental Data 1) was synthetized using the GeneArt service from ThermoFischer Scientific and subsequently cloned into the *Pac*I restriction site into the pKLV-U6-Braf gRNA-PGKpuro2ABFP.

### Cell Lines, Transfections, Infections and Reagents

The mouse embryo fibroblast NIH-3T3-TVA cells, kindly provided by Eric Holland, were cultured in DMEM (Sigma-Aldrich, Cat. D5796) + 10% CS (Sigma-Aldrich, Cat. C8056). The Gp2-293 packaging cell line (Clontech, Cat. 631458) were grown in DMEM (Sigma-Aldrich, Cat. D5796) + 10% FBS (Sigma-Aldrich, Cat. F7524). DF1 chicken fibroblasts (ATCC, Cat. CRL-12203) were grown at 39°C in DMEM containing GlutaMAX-I (Gibco, Cat. 31966-021) and 10% FBS (Sigma-Aldrich, Cat. F7524). The mouse neuronal stem cells (NSCs) used to test gRNA *in vitro* and *in vivo* were derived from the whole brain of newborn mice of *Ntv-a; LSL-Cas9* and *Gtv-a; hGFAP-Cre; LSL-Cas9; p53*^*lox/lox*^, respectively. NSCs and tumorspheres were grown in Mouse NeuroCult proliferation kit (Stem Cell Technologies, Cat. 05702), supplemented with 10ng/ml recombinant human EGF (Gibco, Cat. PHG0313), 20ng/ml basic-FGF (Sigma-Aldrich, Cat. F0291-25UG), and 1mg/ml Heparin (Stem Cell Technologies, Cat. 07980).

*Ntv-a; LSL-Cas9* and NIH-3T3-TVA cells were subsequently infected with either the pMSCVhygro-CRE or lentiCas9-Blast, respectively, to induce Cas9 expression.

DF1 cells were transfected with the different RCAS-gRNA retroviral plasmids using FuGENE 6 Transfection reagent (Promega, Cat. E2691), accordingly to manufacturer's protocol. DF1 RCAS-virus containing media was used to infect NSCs and NIH-3T3-TVA-Cas9. NSCs were infected with four cycles of spin infection (1000rpm for 2hr) and then selected with 1µg/ml Puromycin (Sigma-Aldrich, Cat. P8833-25MG).

Viruses, other than RCAS, were generated in Gp2-293 using calcium-phosphate precipitate transfection: lentiviruses (pKLV-U6gRNA-PGKpuro2ABFP and lentiCas9-Blast) were produced by co-transfection with 2nd generation packaging vectors (pMD2G and psPAX2) and retroviruses (pMSCVhygro-CRE) with VSVg packaging vector. High-titer virus was collected at 36 and 60hr following transfection and used to infect cells in presence of 7µg/ml polybrene (Sigma-Aldrich, Cat. H9268-5G) for 12hr. Transduced cells were selected after 48hr from the last infection with Blasticidin (3µg/ml) (Gibco, Cat. A11139-03) or Hygromycin (300µg/ml) (Sigma-Aldrich, Cat. H3274-25MG).

Entrectinib (RXDX-101) and Dabrafenib (GSK2118436) were purchased from Selleckchem (Cat. S7998 and S2807, respectively).

### NSCs and Tumorspheres Preparation

For the derivation of mouse NSCs and tumor neurospheres, the tissue was enzymatically digested with 5 ml of papain digestion solution (0.94 mg/ml papain (Worthington, Cat. LS003119), 0.48mM EDTA, 0.18mg/ml *N*-acetyl-L-cysteine (Sigma-Aldrich, Cat. A9165-5G) in Earl's Balanced Salt Solution (EBSS) (Gibco, Cat. 14155-08)) and incubated at 37°C for 8min. After digestion, the enzyme was inactivated by the addition of 2ml of 0.71mg/ml ovomucoid (Worthington, Cat. LS003087) and 0.06mg/ml DNaseI (Sigma-Aldrich, Cat. 10104159001) diluted in Mouse Neurocult NSC basal medium (Stem Cell Technologies, Cat. 05700) without growth factors. The cell suspension was then passed through a 40µm mesh filter to remove undigested tissue, washed first with PBS and then with 3 ml of ACK lysing buffer (Gibco, Cat. A1049201). Single cells suspension was then centrifuged at a low speed and resuspended in Mouse NeuroCult proliferation kit (Stem Cell Technologies, Cat. 05702), supplemented with 10ng/ml recombinant human EGF (Gibco, Cat. PHG0313), 20ng/ml basic-FGF (Sigma-Aldrich, Cat. F0291-25UG), and 1mg/ml Heparin (Stem Cell Technologies, Cat. 07980).

### Free Floating ImmunoFluorescence (FF-IF)

Adult (4-6 weeks) and pups (1 day) brains were fixed with PFA 4% (Electron Microscopy Sciences, Cat. 15713) and then incubated with sucrose 15% and 30%. Each step was done overnight at 4°C. Brains were then sectioned by using a sliding microtome with freezing stage (Fisher). Sections of 80 µm were blocked in Goat Serum 10%, BSA 2%, Triton 0.25% and mouse on mouse blocking reagent (Vectors Laboratories, Cat. BMK-2202) in PBS for 2hr at room temperature (RT). Primary antibodies were incubated overnight at 4°C in the blocking solution and the following day for 30 min at RT as detailed: GFAP (Millipore, MAB360, 1:500), NESTIN (BD Pharmingen, #556309, 1:100) and EGFP (Aves Labs, GFP-1010, 1:1000). Slices were then washed in PBS-Triton 0.25% and incubated with the secondary antibody for 2hr. Secondary antibodies were from Invitrogen (Alexa-Fluor anti-chicken^488^, anti-rabbit^555^, anti-mouse^555^). After extensive washing in PBS-Triton 0.25%, nuclei were stained with DAPI for 3 min at RT. Sections were mounted with ProLong Gold Antifade reagent (Invitrogen, Cat. P10144).

Brain mapping was performed with a TCS SP5 confocal microscope (Leica Microsystems) equipped with Leica HCS-A and custom made iMSRC software ^57^. Final images were acquired with a 20x 0.7 N.A. dry objective. The regions of interest definition were done on mosaics of the full brain sections acquired with a 10 x 0.4 N.A. dry objective.

### Immunoblotting

Cell pellets were lysed with JS lysis buffer (50mM HPES, 150mM NaCl, 1% Glycerol, 1% Triton X-100, 1.5mM MgCl_2_, 5mM EGTA) and protein concentrations were determined by DC protein assay kit (Biorad). Proteins were separated on house-made SDS-PAGE gels and transferred to nitrocellulose membrane (Amersham). Membranes were incubated in blocking buffer (5% milk 0.1% Tween, 10 mM Tris at pH 7.6, 100 mM NaCl) and then with primary antibody either 1 hour at room temperature or overnight at 4°C according to the antibody. Antibodies used for western-blot are: TP53 (Cell Signaling Technology, #2524, 1:1000), PTEN (Cell Signaling Technology, #9188, 1:1000), CDKN2A (Santa Cruz Biotechnology, sc-32748, 1:500), pERK (Cell Signaling Technology, #9101, 1:2000), total ERK (Cell Signaling Technology, #9102, 1:1000), pMEK (Cell Signaling Technology, #9154, 1:500), total MEK (Santa Cruz Biotechnology, sc-219, 1:500) and VINCULIN (Sigma-Aldrich, V9131, 1:10000). Anti-mouse or rabbit-HRP conjugated antibodies (Jackson Immunoresearch) were used to detect desired protein by chemiluminescence with ECL (Amersham, RPN2106).

### Immunohistochemistry

Tissue samples were fixed in 10% formalin, paraffin-embedded and cut in 3µm sections, which were mounted in superfrostplus microscope slides and dried. Tissues were deparaffinized in xylene and re-hydrated through a series of graded ethanol until water. For histopathological analysis, sections were stained with hematoxylin and eosin (H&E). For immunohistochemistry, paraffin sections underwent first antigenic exposure process, endogenous peroxidase was blocked and the slides were then incubated in blocking solution (2.5% BSA, 10% goat serum, with or without mouse on mouse IgG (MOM), according to the species of primary antibody, in PBS). Incubation with the appropriate primary antibodies was carried out over-night as detailed: GFAP (Millipore MAB360, 1: 500), NeuN (Millipore, MAB377, 1:100), OLIG2 (Millipore, AB9610, 1:400), NESTIN (BD Pharmingen, #556309, 1:100), PTEN (Cell Signaling Technology, #9559, 1:100) and CDKN2A (Santa Cruz Biotechnology, sc-32748, 1:100). After incubating with the primary antibody, all slides were incubated with appropriate secondary antibodies and the visualization system AB solution (AB solution-Vector, Ref. PK-6100). Finally, slides were dehydrated, cleared and mounted with a permanent mounting medium. The immunohistochemistry for TP53 (CNIO monoclonal antibody core, clone POE316A, 1:100), KI67 (Master Diagnostica, #0003110QD, undiluted), Cas9 (Cell Signaling Technology, #14697, 1:100), pan-TRK (Cell Signaling Technology, #92991, 1:100) were performed using an automated immunostaining platform (Ventana discovery XT, Roche). BRAF^V600E^ immunostaining was performed on a Leica Bond-III stainer (Leica Biosystem, Newcastle, UK) with a 1:100 dilution of anti-BRAF^V600E^ (VE1) mouse monoclonal primary antibody (Spring Bioscience, Pleasanton, CA).

### Blood Counts and Flow Cytometry

For the analysis of the Cas9-induced immune response, 4 weeks old *Ntv-a; LSL-Cas9*; *hUBC-CreERT2* mice were fed *ad libitum* with tamoxifen containing diet for the duration of the experiment (Supplementary Fig. 3a).

Complete blood counts were carried out using the Abacus Junior Vet (Diatron). Cells were isolated from spleen and brain by mechanical disruption. Red Blood Cells were lysed using the red blood cell lysis buffer (Sigma-Aldrich). All cells were stained with CD45-PerCP (Biolegend, #103130, 1:200), CD3-AF700 (eBiosciences, #56-0032, 1:100), CD4-PECy7 (BD Pharmingen, #552775, 1:200), Gr-1-PE (BD Pharmingen, #553128, 1:200), CD11b-PerCPCy5.5 (BD Pharmingen, #550993, 1:30) and B-220-APC-CY7 (BD Pharmingen, #552094, 1:200). Samples were acquired in an LSR Fortessa (BD, San Jose CA) equipped with 355nm, 488nm, 561nm and 640nm lines. We used pulse processing to exclude cell aggregates and DAPI to exclude dead cells. All data were analyzed using FlowJo 9.9.4 (Treestar, Oregon).

### Cell Proliferation, Soft-Agar Assay and Cell Cycle Analysis

NSCs and tumorspheres cells were seeded in 96-well culture plates (4,000 per well) in quintuplicate and treated for 96hr. At the end of the incubation period, survival of cells was determined by the MTT assay. Briefly, MTT was added to each well and samples were incubated for 4h before lysing in formazan dissolving solution. Colorimetric intensity was quantified using an ELISA reader at 590 nm. Values were obtained after subtraction of matched blanks (medium only). The OD values of DMSO controls were taken as 100% and values for drug treatment are expressed as % of control.

The soft-agar growth assay was performed by seeding cells in triplicates at 300,000 cells/well in NSCs culture medium containing 0.4% Noble agar (Sigma-Aldrich, Cat. A5431). Cells were plated on top of a layer of NSCs culture medium containing 0.65% Nobel agar. Colonies were stained 3 weeks after plating with 2mg/ml of thiazolyl blue tetrazolim bromide (Sigma-Aldrich, Cat. M5655) for 1h at 37°C.

For the cell cycle analysis, cells were fixed with 70% cold ethanol for 2hr. Fixed cells were treated with RNAse for 20min before addition of 50µg/ml propidium iodide (PI) and analyzed by FACS.

### Reverse Transcription Quantitative PCR and Analysis of cDNA fragments

RNA from NSCs and frozen tissue was isolated with TRIzol reagent (Invitrogen, Cat. 15596-026) according to the manufacturer’s instructions. For reverse transcription PCR (RT-PCR), 500ng of total RNA was reverse transcribed using the High Capacity cDNA Reverse Transcription Kit (Applied Biosystems, Cat. 4368814). The cDNA was used either for quantitative PCR or Sanger sequencing. Quantitative PCR was performed using the SYBR-Select Master Mix (Applied Biosystems, Cat. 4472908) according to the manufacturer’s instructions. qPCRs were run and the melting curves of the amplified products were used to determine the specificity of the amplification. The threshold cycle number for the genes analyzed was normalized to GAPDH. Sequences of the primers used are listed in (Supplementary Table1).

For Sanger sequencing PCR fragments, cDNA was PCR-amplified using primers listed in (Supplementary Table1), in-gel purified and ligated into the pGEM-T Easy vector (Promega, Cat. A1360) and submitted to sequence.

### Genomic DNA Isolation and Analysis

Genomic DNA was isolated by proteinase K/sodium dodecyl sulfate (SDS)/phenol extraction method described briefly below. Cell pellets were incubated in lysis buffer (10mM Tris-HCl ph8, 100mM NaCl, 0.5mM EDTA, 10% SDS and proteinase K) for 4h at 55°C. Samples were extracted using phenol:chloroform (1:1) and Phase Lock heavy 2ml tubes (5PRIME, Cat. 2302830). The aqueous phase was recovered to fresh tubes and 0.1M sodium acetate and 100% cold ethanol were added. Samples were centrifuged at 15000rpm for 25min. After washing in 70% cold ethanol, draining and dissolving in water, genomic DNA was quantified. 100ng of DNA were amplified with specific primers listed in (Supplementary Table1). PCR products were cloned into the pGEM-T Easy vector and submitted for sequencing.

### Fluorescence *in situ* Hybridization (FISH)

Two sets of FISH probes were used to study the deletion between the *Ntrk1* and *Bcan* mouse genes. BMQ-437D10 bacterial artificial chromosome (BAC) that map at the intergenic *Ntrk1-Bcan* (3qF1 cytoband), was purchased from Source Bioscience and labelled by Nick translation assay with Texas Red fluorochrome to generate a locus-specific FISH probe. BMQ-386N22 BAC clone (3qA3 cytoband) was labelled with Spectrum Green fluorochrome to generate a control probe to enumerate mouse chromosome 3. FISH analyses were performed according to the manufacturers’ instructions, as previously described ^58^, on Carnoy’s fixed cells mounted on positively charged slides (SuperFrost, Thermo Scientific). Briefly, the slides were first dehydrated followed by a denaturing step in the presence of the FISH probe at 85°C for 10min and left overnight for hybridization at 45°C in a DAKO hybridizer machine. Finally, the slides were washed with 20×SSC (saline-sodium citrate) buffer with detergent Tween-20 at 63°C, and mounted in fluorescence mounting medium (DAPI). FISH signals were manually enumerated within nuclei. FISH images were also captured using a CCD camera (Photometrics SenSys camera) connected to a PC running the Zytovision image analysis system (Applied Imaging Ltd., UK) with focus motor and Z stack software.

### Analysis of gene expression in normal and tumor tissues

RNA-seq data for human normal brain samples were downloaded from the GTEx data portal (https://www.gtexportal.org/). ENCODE mouse brain expression data were downloaded from the NCBI (https://www.ncbi.nlm.nih.gov/gene/). RNA-seq data for *NTRK1* in the TCGA GBMLGG dataset were downloaded from the GlioVis data portal (http://gliovis.bioinfo.cnio.es) ^59^. Sample IDs of patients carrying *NTRK1* gene fusions were either previously described ^28^ or obtained from the TCGA Fusion gene Data Portal (http://54.84.12.177/PanCanFusV2/).

### Statistical analysis

Data in bar graphs are presented as mean and SD, except otherwise indicated. Results were analyzed by unpaired two-tailed Student's *t*-tests using the R programming language ^60^. Kaplan–Meier survival curve were produced using the “survminer” R package and *P* values were generated using the Log-Rank statistic. Box-plots were made with the “ggplot2” R package. Drug dose response curves were produced with GraphPad Prism.

### Mouse Strains and Husbandry

*Nestin-tv-a* and *GFAP-tv-a* ^8,9^ were generously provided by Eric Holland. *Rosa26-LSL-Cas9* knockin mouse strain ^4^ was purchased from The Jackson laboratory (Cat. 024857). *Nestin-Cre* ^11^, *hGFAP-Cre* ^10^, *hUBC-CreERT2* ^12^ transgenic lines were kindly provided by various researchers at the Spanish National Cancer Research Center (Marcos Malumbres, Mariano Barbacid and Maria Blasco).

Mice were housed in the specific pathogen-free animal house of the Spanish National Cancer Centre under conditions in accordance with the recommendations of the Federation of European Laboratory Animal Science Associations (FELASA). All animal experiments were approved by the Ethical Committee (CEIyBA) and performed in accordance with the guidelines stated in the International Guiding Principles for Biomedical Research Involving Animals, developed by the Council for International Organizations of Medical Sciences (CIOMS).

### Generation of Murine Gliomas

For the RCAS-mediated gliomagenesis, newborns or 4-6 weeks old mice, were injected intracranially with 4x10^5^ DF1 cells 1:1 dilution between RCAS-PDGFB and RCAS-gRNA expressing cells per mouse. For the *p53-null* TVA-Cas9 NSCs infected with the RCAS-gRNA-pairs or the pKLV-Braf-V637E-HDR, 4-5 weeks old immunodeficient *NOD/*SCID mice were injected intracranially with 5 x10^5^ cells. Adults mice were anaesthetized by 4% isofluorane and then injected with a stereotactic apparatus (Stoelting) as previously described ^13^.

For the Cas9-inducible tumor model (*Ntv-a; LSL-Cas9*; *hUBC-CreERT2),* two weeks after DF1 RCAS-gRNA plasmid injection, mice received intraperitoneal injections of 4-Hydroxytamoxifen (Sigma-Aldrich, Cat. H6278) (2mg/injection, 4-6 injections).

After intracranial injection, mice were checked until they developed symptoms of disease (lethargy, poor grooming, weight loss, macrocephaly).

## Author contributions

BO supervised and performed experiments and contributed to write the manuscript. AC-G, CM and VM performed experiments. OU contributed to the analysis of the immune response. SR-P and RR-T performed the FISH analysis. JH helped with histopathological analysis of the tumor tissues. MS designed, supervised and performed experiments and wrote the manuscript.

## Acknowledgments

ACG is recipient of a Severo-Ochoa PhD fellowship. CM and VM are recipient of a “La Caixa” PhD fellowship. We thank A. J. Schuhmacher for the initial assistance with the intracranially injections in adult mice and C.S. Clemente-Troncone for the technical support. We thank David Olmeda and Marisol Soengas for sharing reagents. We sincerely thank Dr. José Luis Rguez Peralto (“Hospital U. 12 de Octubre”, Madrid) for the BRAF V600 IHCs staining. This research was supported by funds from the “Acción Estratégica en Salud” Spanish National Research and Development Plan, Instituto de Salud Carlos III (ISCIII), cofounder by FEDER (ERDF) (PI14/01884) to S.R-P, by a “*Beca Leonardo a Investigadores y Creadores Culturales 2017*” from the BBVA Foundation and a grant from the Seve Ballesteros Foundation to MS.

